# Synthetic Cooling Agents in US-marketed E-cigarette Refill Liquids and Popular Disposable E-cigarettes: Chemical Analysis and Risk Assessment

**DOI:** 10.1101/2021.06.09.446946

**Authors:** Sairam V. Jabba, Hanno C. Erythropel, Deyri Garcia Torres, Lauren A. Delgado, Jackson G. Woodrow, Paul T. Anastas, Julie B. Zimmerman, Sven-Eric Jordt

## Abstract

**Background:** Menthol, through its cooling and pleasant sensory effects, facilitates smoking and tobacco product initiation, resulting in the high popularity of mint/menthol-flavored E-cigarettes. More recently, E-cigarette vendors started marketing synthetic cooling agents as additives that impart a cooling effect but lack a characteristic minty odor. Knowledge about content of synthetic coolants in US-marketed E-cigarette products and associated health risks is limited.

**Methods:** E-liquid vendor sites were searched with the terms “koolada”, “kool/cool”, “ice”, or WS-3/WS-23, denoting individual cooling agents, and relevant refill E-liquids were purchased. “Ice” flavor varieties of Puffbar, the most popular disposable E-cigarette brand, were compared with non-”Ice” varieties. E-liquids were characterized, and synthetic coolants quantified using GC/MS. Margin of exposure (MOE), a risk assessment parameter, was calculated to assess the risk associated with synthetic coolant exposure from E-cigarette use.

**Results:** WS-3 was detected in 24/25 refill E-liquids analyzed. All Puffbar flavor varieties contained either WS-23 (13/14) or WS-3 (5/14), in both “Ice”- and non-”Ice” flavors. Modeling consumption of WS-3 from vaped E-liquids, resulted in MOEs below the safe margin of 100 for most daily use scenarios. MOEs for WS-23 were <100 for 10/13 Puffbar flavors in all use scenarios. Puffbar power specifications are identical to Juul devices.

**Conclusions:** Synthetic cooling agents (WS-3/WS-23) were present in US-marketed E-cigarettes, at levels that may result in consumer exposures exceeding safety thresholds set by regulatory agencies. Synthetic coolants are not only found in mint-or menthol-flavored products, but also in fruit- and candy-flavored products, including popular disposable E-cigarette products such as Puffbar.

**Implications:** Synthetic cooling agents are widely used in “kool/cool”- and “ice”-flavored E-liquids and in E-liquids without these labels, both as a potential replacement for menthol or to add cooling ‘notes’ to non-menthol flavors. These agents may be used to bypass current and future regulatory limits on menthol content in tobacco products, and not just E-cigarettes. Since synthetic cooling agents are odorless, they may not fall under the category of “characterizing flavor”, potentially circumventing regulatory measures based on this concept. Regulators need to consider the additional health risks associated with exposure to synthetic cooling agents.

## Introduction

Additives and flavors are added to tobacco products to counteract the harshness, sensory irritation and bitter tastes of tobacco and nicotine. In the United States, the Family Smoking Prevention and Tobacco Control Act (FSPTCA) of 2009 authorized the Food and Drug Administration (FDA) to regulate constituents, additives, and flavors in tobacco products ^1^. The FSPTCA banned the sales of flavored combustible cigarettes, with the exception of menthol cigarettes ^1^. Menthol in tobacco products is well understood to reduce harshness, impart a pleasant minty and cooling flavor and increase palatability ^2^. Menthol affects tobacco and nicotine use behaviors such as initiation, dependence, abuse liability and reduces the ability to quit, especially among the youth and young adults ^2^. The use of mentholated tobacco products is associated with tobacco use disparities, with 85% of African American smokers using menthol cigarettes ^2,3^. The US FDA announced in May 2021 that it will issue a rule to ban menthol cigarettes ^4^. Menthol is also a popular flavor additive in E-cigarettes, especially after the FDA banned all flavored closed pod systems (eg. Juul) except for menthol and tobacco ^5^. Sales of menthol Juul pods increased 6-fold from August 2019 to May 2020 (10.7% to 61.8%) when all other Juul flavors (besides tobacco) were removed from the marketplace ^6,7^.

Synthetic cooling agents share the sensory cooling effects of menthol, but lack menthol’s strong minty odor, especially at the concentrations at which they are added to E-liquids ^8,9^. These agents were developed beginning in the 1970s, to replace menthol in body care and food products. In the 1970s and 80s, the tobacco industry including R.J. Reynolds and Phillip Morris, carried out consumer tests with cigarettes containing synthetic cooling agents, to impart a “cooling without menthol” experience ^8–11^. A 2018 analysis of combustible cigarettes (both menthol and non-menthol) marketed in Germany detected WS-3, indicating that coolants other than menthol are added to tobacco products ^12, 13^. In a recent study we demonstrated that European-marketed Juul menthol pods contained the synthetic cooling agent WS-3, while US and Canadian pods did not ^14^. More recently Omaiye et al., demonstrated the presence of synthetic coolants in US-marketed disposables ^15^. While synthetic cooling agents are sold to (<u>D</u>o-<u>I</u>t-<u>Y</u>ourself) DIY users in the US, often under the label “Koolada”, the identity and concentrations of these agents in US-marketed E-cigarettes and E-cigarette refill liquids (“E-liquids”) remains unknown ^16^.

While natural and synthetic mint- and cooling compound are widely used in food and other consumer products, there are rising concerns about their toxicity, especially in poorly regulated products such as E-cigarettes. For example, we detected concerning levels of pulegone, a carcinogenic mint flavorant banned by FDA in food, in a variety of mint- and menthol-flavored E-cigarette and smokeless tobacco products ^17,18^. Synthetic cooling agents such as WS-3 are classified by the Flavor Extracts Manufacturers Associaton (FEMA) as GRAS (Generally Recognized As Safe). However, this applies only to their intended use in food products, but not to E-cigarette use resulting in inhalational exposure ^19^. The Joint Expert Committee on Food Additives (JECFA) of the World Health Organization (WHO) established a threshold of concern of 90 μg/day/person for intake of WS-3 and related compounds, a level that is often exceeded by consumers ^20^. Based on oral toxicity studies, an oral No-Observed-Adverse-Effect-Level (NOAEL) of 8 mg/kg-bw for WS-3 ^20–22^ was established, and 5 mg/kg-bw for WS-23, another widely used synthetic cooling agent. Chronic or higher doses were observed to cause kidney and liver lesions ^20,23^. *In vitro* genotoxicity studies in mammalian cells demonstrated WS-23 to be clastogenic (i.e., inducing disruption or breakages of chromosomes), leading to a call by JECFA for additional toxicological studies to further evaluate the safety of WS-23 ^20^. Exposure of airway-epithelial cells in-vitro to various concentrations of synthetic coolants and WS-23 containing e-liquids also demonstrated a dose-dependent cellular toxicity ^15^. A recent inhalation exposure study in rodents by an E-liquid manufacturer for WS-23 determined a NOAEL of 29 mg/kg-bw (342 mg/m3) in rodents ^24^. Due to their evident toxicity and increased use in E-cigarettes, it is critical to evaluate the potential health risks associated with synthetic coolant exposure from E-cigarette use.

In the present study, we determined the levels of synthetic cooling agents (WS-3, WS-23) as well as important peppermint-(menthol and menthone) and spearmint-(carvone) flavor chemicals, in US-marketed refill E-liquids and in disposable E-cigarettes of the brand Puffbar. The rationale to include menthol, menthone and carvone in chemical analysis is that, they are the most frequent mint flavors encountered in E-liquids. Several recent E-cigarette prevalence studies indicated high use of disposables, especially ‘ice’-flavored, among youth and young adults ^6,7,25–29^. Disposable E-cigarette brand, Puffbar, was selected because, according to the 2021 US National Youth Tobacco Survey it was determined as the most popular E-cigarette device among middle and high school students ^6,7,28^. In addition, the power specifications for Puffbar devices were determined. We then assessed the health risk associated with synthetic coolant exposure via E-cigarette use by calculating the Margin of Exposure (MOE), a risk assessment parameter, for a range of product use scenarios modeling low, moderate and frequent E-cigarette use.

## Materials and Methods

### Analysis of synthetic cooling agents and other flavor content in US-marketed E-cigarette liquids and their aerosols

#### E-liquid selection

E-cigarette vendor websites keyword searches for “koolada”, “kool”, “WS-3”, and “WS-23” were carried out to identify E-liquids potentially containing synthetic coolants. Identified products, including a concentrate (“Agent Cool”) for do-it-yourself (DIY) users, were purchased directly from vendors in 2019 and 2020. In addition, previously purchased menthol/mint/spearmint/wintergreen E-liquids (2019) were re-analyzed for synthetic coolant presence. In early 2020, all available flavors of “Puffbar” were purchased from puffecig.com due to the frequent occurrence of the words “cool/ice” in Puffbar flavor names (see Table 1,2 for an overview). Purchased E-liquids were stored in dark at room temperature.

**Table 1:**
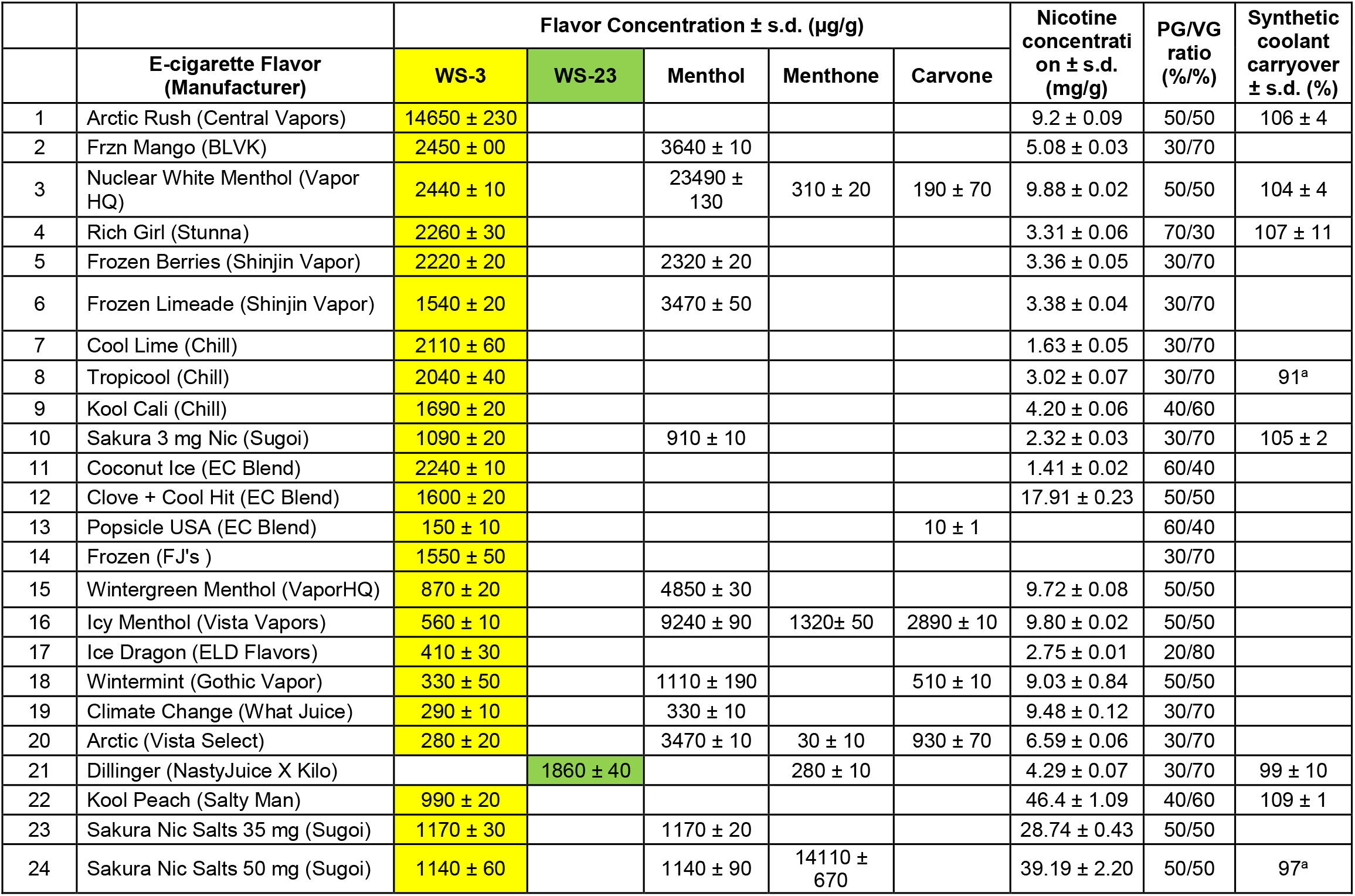
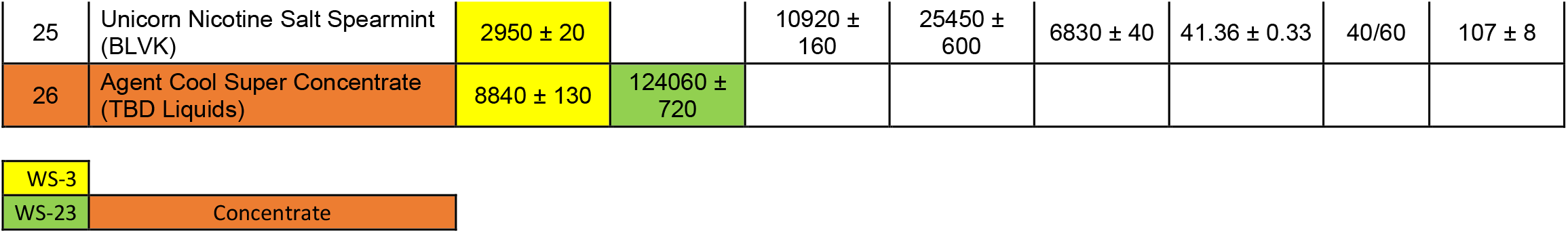
Concentrations of synthetic cooling agents (WS-3 and WS-23), menthol, menthone, carvone, nicotine and solvent chemicals in US-marketed E-cigarette refill liquids. Carryover rates for synthetic coolants are provided for select refill liquids. ^a^single experiment. WS-3 and WS-23 concentrations are highlighted in yellow and green, respectively; Concentrate E-liquid highlighted in orange.

**Table 2:**
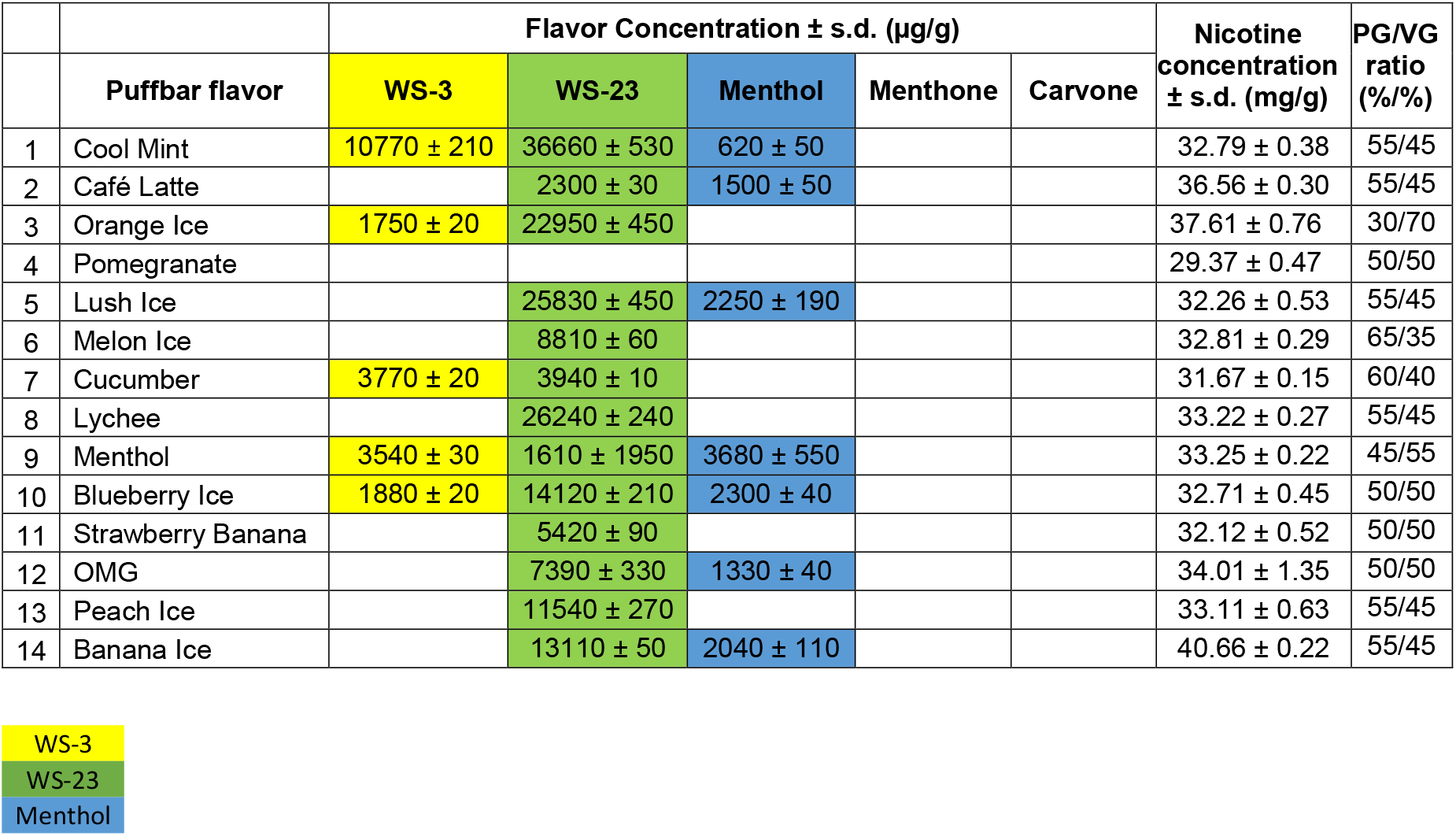
Chemical concentrations of synthetic cooling agents (WS-3 and WS-23), menthol and nicotine in flavors of US-marketed Puffbar disposable E-cigarettes. No menthone or carvone was found in any of the analyzed Puffbar e-liquids. PG/VG ratio measured in triplicate and rounded to next 5. Empty cells represent “not detected”.

| | Puffbar flavor | Flavor Concentration ± s.d. ($\mu\text{g/g}$) | | | | | Nicotine concentration ± s.d. (mg/g) | PG/VG ratio (%/%) |
| --- | --- | --- | --- | --- | --- | --- | --- | --- |
|  |  | WS-3 | WS-23 | Menthol | Menthone | Carvone |  |  |
| 1 | Cool Mint | 10770 ± 210 | 36660 ± 530 | 620 ± 50 | | | 32.79 ± 0.38 | 55/45 |
| 2 | Café Latte | | 2300 ± 30 | 1500 ± 50 | | | 36.56 ± 0.30 | 55/45 |
| 3 | Orange Ice | 1750 ± 20 | 22950 ± 450 | | | | 37.61 ± 0.76 | 30/70 |
| 4 | Pomegranate | | | | | | 29.37 ± 0.47 | 50/50 |
| 5 | Lush Ice | | 25830 ± 450 | 2250 ± 190 | | | 32.26 ± 0.53 | 55/45 |
| 6 | Melon Ice | | 8810 ± 60 | | | | 32.81 ± 0.29 | 65/35 |
| 7 | Cucumber | 3770 ± 20 | 3940 ± 10 | | | | 31.67 ± 0.15 | 60/40 |
| 8 | Lychee | | 26240 ± 240 | | | | 33.22 ± 0.27 | 55/45 |
| 9 | Menthol | 3540 ± 30 | 1610 ± 1950 | 3680 ± 550 | | | 33.25 ± 0.22 | 45/55 |
| 10 | Blueberry Ice | 1880 ± 20 | 14120 ± 210 | 2300 ± 40 | | | 32.71 ± 0.45 | 50/50 |
| 11 | Strawberry Banana | | 5420 ± 90 | | | | 32.12 ± 0.52 | 50/50 |
| 12 | OMG | | 7390 ± 330 | 1330 ± 40 | | | 34.01 ± 1.35 | 50/50 |
| 13 | Peach Ice | | 11540 ± 270 | | | | 33.11 ± 0.63 | 55/45 |
| 14 | Banana Ice | | 13110 ± 50 | 2040 ± 110 | | | 40.66 ± 0.22 | 55/45 |

WS-3
WS-23
Menthol

#### Power specifications measurement

Puffbar devices were opened manually, and the heating coil was carefully pulled towards the front of the opened device by the connecting wires. Resistance was measured in three devices by clamping a digital multimeter (Fluke 116) to the connecting wires. Battery specifications were printed onto the battery pack and reported as such.

#### E-liquid chemical analysis

E-liquids were analyzed using an established GC-MS method and selected compounds (WS-3, WS-23, menthol, menthone, carvone, nicotine) were quantified in triplicate using an established GC-FID method ^30^. To do so, commercially available standards of N-Ethyl-p-menthane-3-carboxamide (WS-3; 99%; CAS# 39711-79-0), (S)-nicotine (>99%; CAS# 54-11-5), (DL)-menthol (99%; CAS# 89-78-1), (D)-carvone (>96%; CAS# 2244-16-8; all Sigma-Aldrich, Saint Louis, MO), 2-isopropyl-N,2,3-trimethylbutyramide, (WS-23; >98%; CAS# 51115-67-4, TCI, Portland, OR), (L)-menthone (97%; CAS# 14073-97-3; Alfa Aesar, Haverhill, MA), propylene glycol (>99.5%; CAS# 57-55-6), and glycerol (99.6%; CAS 56-81-5; both Fisher Scientific, Hampton, NH) were used to construct calibration curves in the relevant concentration range. Samples were diluted in 1mL of methanol [JT Baker, Center Valley, PA] containing 1 g/L of 1,4-dioxane [Alfa Aesar] as internal standard (IS) and injected; the detailed analytical methods have been described previously ^30^. PG/VG ratios for Puffbars were rounded to the nearest 5 for simplicity.

#### Coolant carryover analysis

To assess the carryover of the synthetic coolants WS-3 and WS-23 from refill E-liquids to E-cigarette-generated aerosol, the Suorin iShare refillable pod E-cigarette device was used as its power specifications are very similar to the Puffbar and Juul device (Suorin iShare: 3.7V, 1.8Ω, resulting max. power 7.6W; Puffbar and Juul: 3.7V, 1.7Ω, resulting max. power 8.1W). Moreover, Suorin remains highly popular among vapers, and especially high school age vapers, as a favorite refillable device ^29^. The Sourin device was attached via a custom-3D printed connector to a previously described house-built vaping machine ^31^. In brief, the device was controlled by a programmable Arduino board that activates a micro-diaphragm pump at an operator-defined puff length, inter-puff break, and number of puffs. A microneedle valve in combination with a flow meter (Omega, Norwalk, CT) was used to control the flow rate. Generated aerosol was trapped by a series of liquid nitrogen-chilled cold-finger traps. The carryover puffing regime was 20 puffs, 2L/min flow rate, 2.4s puff length, 80mL puff volume, 30s inter-puff break and carryover was determined in triplicate (three separate cartridges). The chosen puffing regime is at the lower spectrum of observed user behavior ^31,32^ and has been utilized previously to determine carryover in popular Juul E-cigarettes ^33^. After 20 puffs were collected for each experiment, the traps were allowed to thaw and the captured material was taken up in 1mL of methanol containing IS and injected into the GC/FID. Percent carryover was calculated as coolant content in trapped aerosol over coolant content of consumed liquid, as determined by weighing the e-liquid-containing cartridge before and after the carryover experiment using an analytical balance (Sartorius Practum, Göttingen, Germany). E-liquids for carryover experiments were chosen based on type of synthetic coolant present (WS-3, WS-23), and based on high, average, or low synthetic coolant and/or nicotine content (Table1).

##### Margin of Exposure (MOE) calculation for synthetic cooling agents

MOE, a preferred risk assessment parameter for toxic compounds by the JECFA ^34^,^35^, was calculated for WS-3 and WS-23 containing E-liquids using the following formula, as described previously:^36^

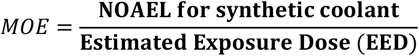

Where NOAEL is the No-Observed-Adverse-Effect-Level, a dose determined as the threshold producing a toxic effect in animal oral toxicity studies. Though <u>b</u>ench<u>m</u>ark <u>d</u>ose (BMD) is the preferred toxicological threshold for MOE calculations, NOAEL-values are applicable when available toxicity data is not amenable for statistical modeling of BMD ^37^, as was the case for WS-3 and WS-23. JECFA determined NOAELs for WS-3 and WS-23 are 8 and 5 mg/kg-bw/day, respectively ^21–23^. Daily <u>E</u>stimated <u>E</u>xposure <u>D</u>oses were determined from WS-3 and WS-23 concentrations in the E-liquids and their estimated average daily amount consumed. Nicotine concentrations in the majority of refill E-liquids ranged between 0.1-1.0%, and user surveys found consumption volumes for such low nicotine E-cigarette liquids to be 9 mL/day, ranging from 1.7-17.7 mL/day) ^18,38,39^. To cover a wide range of user behaviours, MOEs for these refill E-liquids were calculated for consumption volumes of 1-,3-,5-,10- and 15-mL/day. For nicotine salt-containing refill liquids (nicotine content >3%), MOEs were calculated for 1-,3-, 5-mL/day to account for the higher nicotine content. Puffbar E-liquid MOEs were calculated for consumption amounts of half-,1- and 2-Puffbar(s) to model volumes consumed by light, moderate, and heavy users of 3-5% nicotine salt-containing disposable E-cigarettes (Puffbar volume is 1.3 mL). For the calculations, a user body weight of 60 kg was assumed; to convert E-cigarette liquid consumption from volume in mL/day to g/day, the densities of propylene glycol and glycerol (1.04 and 1.26 g/cm^3^, respectively) were taken into consideration.

## Results

### Chemical analysis of synthetic cooling agents, nicotine and other flavorants in US-marketed E-cigarette refill liquids, concentrates and their aerosols

Using our search criteria, 25 E-cigarette refill liquids and 1 concentrate from 19 different brands were identified and their contents of synthetic coolants, mint-related compounds and nicotine identified by GC/MS and quantified by GC/FID. Of the analyzed liquids, 21 contained free-base nicotine and 4 nicotine salts. WS-3 was present in 24/25 E-liquids, with amounts varying from 150 μg/g-14,650 μg/g (Table1). One E-liquid contained WS-23 exclusively (Dillinger;1,860 μg/g)(Table1). The concentrate contained both WS-3 (8,840 μg/g) and WS-23 (124,060 μg/g)(Table1).

Carryover rates for WS-3 from 8-different E-liquids into aerosol ranged between 91-109% (vs. neat E-liquid content) (average: 105±6.4%; n=20; Table1). For the only refill E-liquid it was detected in, the carryover rate for WS-23 was 99±10% (vs. neat E-liquid content; n=3; Table1).

In addition to WS-3 and WS-23, menthol, menthone, carvone, and nicotine were also quantified, which were detected in 13/25 E-liquids for menthol (330-23,490 μg/g), 6/25 for menthone (30-25,450 μg/g) and carvone (10-6830 μg/g), and 4/25 for menthol, menthone, carvone, WS-3 combined (4,940-46,150 μg/g; see Table1 for details). Nicotine concentrations in 20/21 free-base nicotine containing E-liquids were below 1% (1.41-9.97 mg/g), except for Clove+Cool Hit (17.91 mg/g). Nicotine contents in nicotine-salts containing E-liquids ranged from 28.74-44.6 mg/g (Table1). Carryover rates for menthol, menthone and carvone were calculated from vaping of select refill E-liquids and ranged from 103-107%, 85-106% and 97-105%, respectively (Supplementary Table1). Taken together, these results suggest that synthetic coolants are added to US-marketed E-liquids of a wide range of flavors (Table1), including to those already containing the “natural” coolant menthol. Indicated PG/VG ratios for refill E-liquids in Table 1 represent label information.

### Chemical analysis of synthetic cooling agents and menthol in US-marketed disposable Puffbar E-cigarettes

Several Puffbar flavors are marketed as ‘Ice’ flavors, indicating the presence of a cooling agent. WS-23 was detected in 13/14 Puffbar flavors (1,610-36,660 μg/g), WS-3 in 5/14 flavors (1,880-10,770 μg/g) and menthol in 7/14 analyzed flavors (620-3,680 μg/g; Table2). 3/14 Puffbar flavors contained all three compounds, with a combined concentrations of 8,830-48,050 μg/g (Table2). The nicotine concentrations in the Puffbar E-liquids ranged from 29.4-40.7 mg/g. As no PG/VG ratios were indicated on the product label, ratios were measured by GC/FID (Table 2).

### Puffbar power specifications

Contrary to pod-based E-cigarettes, the disposable puffbar device does not contain a separate “pod” or cartridge that can be exchanged, but the e-liquid is housed within the metal frame of the device, where it is stored in an absorbent material much like in the first generation “cig-a-likes”^31^. The coil resistance was measured at 1.7Ω and the battery indicated a maximum power output of 3.7V, which means the device has the exact power specifications as the Juul device ^14^, in addition to its look that also seem to mimic the Juul device. Maximum power output is thus calculated as 8.05W.

### Margin of Exposure calculations for synthetic coolants in US-marketed refill E-liquids and disposable E-cigarettes

To assess the risk associated with daily intake of WS-3 and WS-23 from E-cigarette use, we calculated the MOEs for different use scenarios (see methods for details). For organ toxicity, a safety margin MOE of >100 is considered low risk, while MOEs <100 require prioritization by regulatory agencies for risk mitigation for a given additive ^37,40^

For the E-liquid with the highest WS-3 concentration (Arctic Rush), MOEs ranged from 28 (1-mL/day) to 2 (15-mL/day;Table S2 and Figure1), and are thereby significantly lower than the recommended safety margin of 100. For consumption volumes of 3-mL/day and 5-mL/day, 13/24 and 18/24 respectively, of WS-3 containing refill E-liquids had a MOE of <100. For 10- and 15-mL/day, MOEs were calculated only for the 20 free-base nicotine containing E-liquids: For 10-mL/day and 15-mL/day consumption volumes, 16/20 and 19/20 WS-3 containing E-liquids respectively, yielded a MOE <100 (Table S2 and Figure1). MOEs for the one WS-23-containing refill E-liquid was <100 in all assessed consumption scenarios, except for 1-mL/day consumption (Table S2 and Figure1). The tested concentrate (“Agent Cool”) contained both WS-3 and WS-23. Based on vendor recommendations to add 1-2 drops of concentrate per pod, we conservatively estimated MOEs for a resulting dilution range of 1-3%. Resulting MOEs for WS-23 when added at 1-3% to E-liquids were <100 at all consumption rates, except for the lowest consumption rate (1-mL/day) at 1% dilution. Taken together, these results indicate that users of synthetic coolants flavored E-liquids are exposed to WS-3 and WS-23 levels that can potentially pose a health risk upon long-term use.

**Figure 1:**
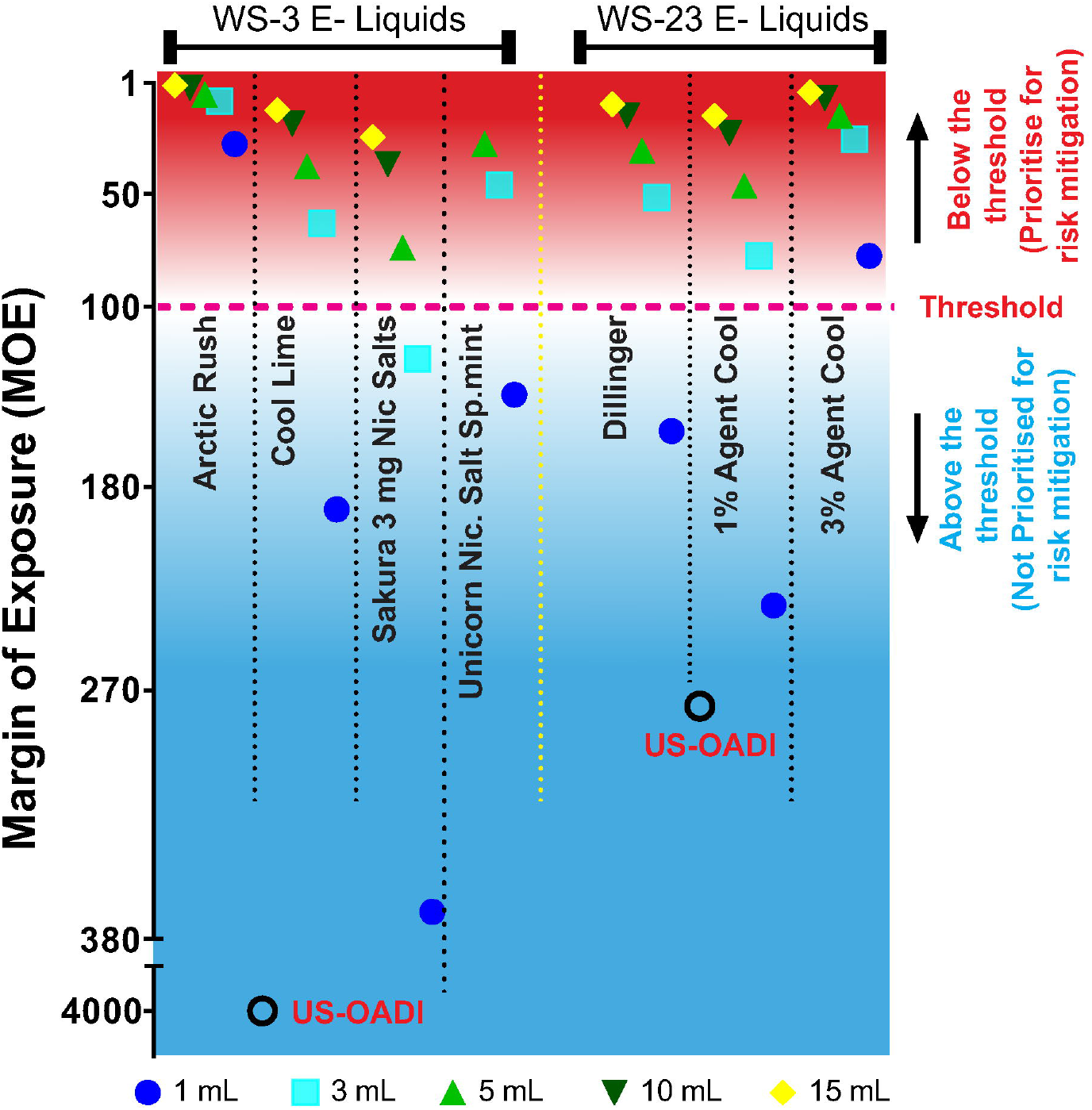
Margins of Exposure (MOE) for synthetic cooling agents from E-Cigarette refill liquids and coolant concentrate. MOEs are plotted for representative WS-3 (left) and WS-23 (right) -containing E-liquids and diluents with high, intermediate and low coolant concentrations, for daily consumption of 1, 3, 5, 10 and 15 ml E-liquid. No-Observed-Adverse-Effect-Levels (NOAEL) determined by JECFA in animal studies of organ toxicity upon oral administration were used for MOE calculations. A MOE of <100 (dashed line) signals a safety concern and requires regulatory mitigation. For WS-23 concentrate (Agent Cool, TBD Liquids), MOEs were calculated for dilutions within the range of manufacturer-recommendations to 1 and 3%. MOEs of the <u>o</u>ral <u>a</u>verage <u>d</u>aily <u>i</u>ntake of US consumers (US-OADI) determined by JECFA are displayed for comparison.

Next, we calculated the MOEs for synthetic coolants for daily consumption of Puffbar devices. For the Puffbar flavor with the highest content of both WS-3 and WS-23 (Cool Mint), the MOEs for consumption of 0.5-2 Puffbars/day ranged from 11-2 for WS-23 and 57-14 for WS-3, values that are significantly lower than the safety margin threshold of 100 (Table S3 and Figure2). MOEs for WS-3 were lower than the safety threshold of 100 at the highest estimated consumption rate (2-Puffbars/day) for all five WS-3 containing Puffbar flavors. MOEs for WS-3 were also <100 for Cucumber and Menthol flavors at 1-Puffbar/day consumption rate. For all other flavors, WS-23 MOEs were <100 for all use scenarios, with the exception of Café Latte (0.5 Puffbar/day), Cucumber (0.5 Puffbar/day) and Menthol (0.5 and 1-Puffbar/day). These results indicate that users of Puffbar disposable E-cigarettes are potentially exposed to levels of synthetic coolants that may pose organ toxicity risks, especially from WS-23 exposure.

**Figure 2:**
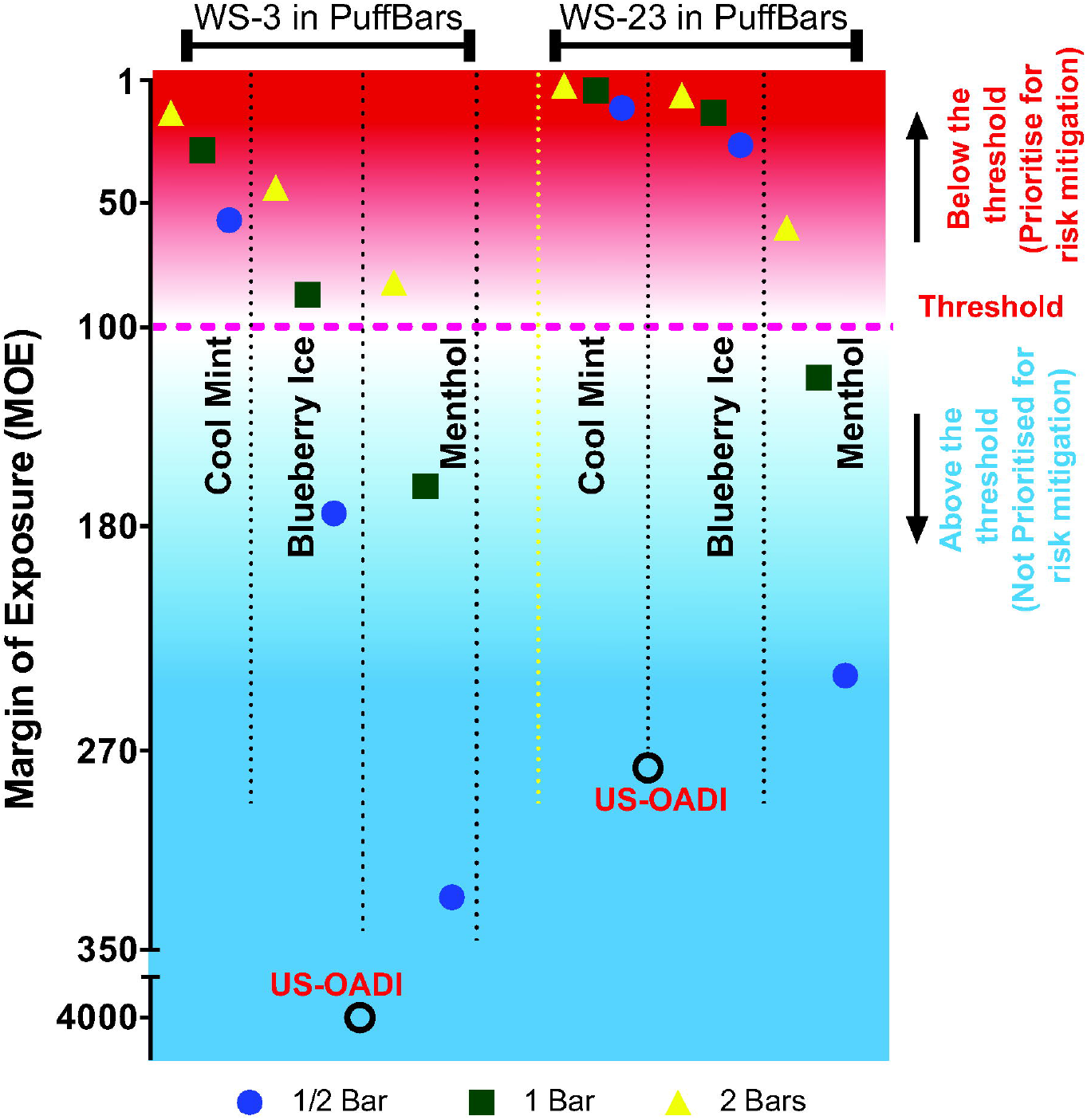
Margins of Exposure (MOE) for synthetic cooling agents from Puffbar-branded disposable E-cigarettes. MOEs are plotted for exposures from Puffbar devices with flavored E-liquids containing both WS-3 and WS-23, at high (Cool mint), intermediate (Blueberry ice) or low (Menthol) concentrations, separated for each coolant (left: WS-3, right: WS-23), calculated for daily consumption of ½ Puffbar (0.65 mL), 1 Puffbar (1.3 mL) and 2 bars (2.6 mL). JECFA-determined NOAEL values were used for MOE calculations. MOEs of the <u>o</u>ral <u>a</u>verage <u>d</u>aily intake of US consumers (US-OADI) determined by JECFA are displayed for comparison.

## Discussion

We detected the presence of synthetic coolants (WS-3 and WS-23) in US-marketed E-cigarette refill liquids and almost all tested flavors of the popular disposable E-cigarette brand, Puffbar, which was found to be the most popular device among US middle- and high school students in 2021 ^28^. Using a relatively low-power E-cigarette that resembles the power specifications of Puffbar and the popular Juul device, we demonstrate that these coolants can efficiently be aerosolized and carried over to reach user airways. Margins of Exposure (MOE) were calculated to assess the health risks associated with exposure to these coolants from E-cigarette use, a parameter used by the US FDA and other global regulatory agencies to regulate the safe use of food additives with carcinogenic and organotoxic effects. These calculations suggests that daily intake of WS-3 or WS-23 by E-cigarette users exceed levels considered acceptable for consumption by FDA and the European Flavour Association (EFFA).

Natural cooling agents such as menthol provide a cooling sensation by activating cold-sensitive trigeminal nerve fibers innervating the upper respiratory tract and oral cavity. These sensory neurons express TRPM8 ion channels, the receptor for menthol that is also activated by cooling ^41,42^. Menthol also has counterirritant, antitussive and analgesic properties and inhibits respiratory irritation responses to tobacco smoke and its various irritant constituents through a TRPM8 dependent mechanism in rodent models ^43,44^. Menthol, through its actions on TRPM8, attenuated the aversive oral effects of nicotine in mice and increased nicotine intake ^45^. The half-maximal concentrations of menthol to activate human TRPM8 is 4μM ^46,47^. This concentration is known to be exceeded after inhalation of menthol cigarette smoke ^48^. A menthol concentration of 0.65μM (16 ppm), was sufficient to reduce sensory irritation by cigarette smoke exposure in mice, illustrating the strong counterirritant properties of menthol even at low concentrations ^43^. In E-cigarette users, menthol is transferred to the airways with high efficiency, where menthol concentrations per puff are comparable to concentrations in puffs generated from menthol cigarettes ^14,33^. Previous analytical studies demonstrated that menthol concentrations in marketed E-liquids range from 2-68,000 μg/mL (13 μM-435 mM) ^49^. In the present study, menthol levels in E-cigarette refill liquids and Puffbar devices ranged from 330 – 23,490 μg/g (~2.4-173 mM) and 620-3,680 μg/g (~4.6-27 mM), respectively. These levels are sufficient to reduce sensory perception of irritation and harshness elicited by nicotine in humans ^50^. Similar to menthol, synthetic cooling agents (including WS-3 and WS-23) also activate TRPM8 with increased efficacy (WS-3) and with comparable half-maximal concentrations (WS-3: 2-4μM; WS-23: 44μM) ^46,51,52^. Several of the refill E-liquids analyzed in this study contained WS-3 at levels comparable to menthol levels in mint- and menthol-flavored E-liquids (150-14,650 μg/g; ~816 μM-80 mM), with some containing synthetic coolants in addition to “natural” cooling agents. The concentrations of WS-3 in several Puffbar E-liquids, up to 1% (1,880-10,770 μg/g; 10-59 mM), are similar to menthol concentrations in liquids of Juul, the market-leading pod E-cigarette system in the US ^14,33^. In addition, the majority of the Puffbar E-liquids analyzed in this study contained WS-23 ranging between 1-3.7% (up to 250 mM). Carryover of synthetic cooling agents from E-liquids into the vapor was highly efficient, approximating 100%, as previously demonstrated for menthol (Table1, SI Table1) ^33^. Based on the puffing regimen in our study, the aerosol concentrations of WS-3 for majority of refill E-liquids (18/24 E-liquids) was calculated to be higher than 5 ppm (range: 5.8-98 ppm). Similarly, WS-3 and WS-23 concentrations for all of disposable Puffbars tested exceeded 10 ppm (WS-3: 12-72 ppm; WS-23: 13-303 ppm), which is sufficient to mediate TRPM8-dependent counterirritant effects. Since these synthetic coolants are either more potent or comparable TRPM8 agonists to menthol, it is likely they facilitate inhalation in vapers using the products analyzed by us. One recent study also quantified menthol and the synthetic coolants (WS-3 and WS-23) in Puffbar devices ^15^. This study (Omaiye et al.,) found similar quantities of synthetic coolants and a higher quantity of menthol in the Puffbar “Cool Mint” product, and higher quantities of all 3 compounds in the “Menthol” product ^15^. Some of these differences in amounts could be that, the tested “Cool Mint” device in this study was smaller in size (1.3 mL) than what was analyzed in Omaiye et al., study (3.2 mL). It is thus possible that new products were introduced and that e-liquid contents might have been adjusted simultaneously. Another observation to explain this difference could be the degree of e-liquid content variability (i.e., production inconsistency and manufacturing sites) of the Puffbar devices, such as evident in WS-23 contents in the “Menthol” product in this study and menthol content in the “Cool Mint” product reported earlier ^15^. This clearly illustrates the variability and inconsistencies of the manufacturing practices in the market place.

Our risk assessment analysis suggests that users of E-cigarettes containing synthetic cooling agents are exposed to amounts of WS-3 and WS-23 that exceed the amounts considered acceptable by regulators for intake from food. Detected amounts were much higher than the thresholds set by JECFA for human intake of flavor additives of this particular class of chemicals (aliphatic and aromatic amines and amides, FEMA Nos 1594-1601, 1606 and 1613) at 90 μg/person/day, either added as individual flavoring or combined total ^20^. Even consumption of just 1-mL of WS-3 containing E-cigarette liquid (150-14650 μg/g) results in exposures exceeding JECFA’s safety threshold by ~2-180 fold. Vaping of 1 disposable Puffbar E-cigarette containing synthetic cooling agents would expose a user to amounts of WS-3 (1,880-10770 μg/g) that exceed the threshold of concern by ~29-176 fold, or ~26-598 fold for WS-23 (3,940-36660 μg/g). There were 5 Puffbar varieties that contained both WS-3 and WS-23, and when added together, would expose a user vaping 1 Puffbar device to ~126-774 fold the recommended maximal daily amount for amides, based on JECFA recommendations. These results strongly suggest that E-cigarette users are exposed to synthetic coolant amounts which would be considered unsafe when consumed in food, raising concerns for health risks upon chronic exposure.

The available chemical toxicity margins for WS-3 and WS-23, such as provided by JECFA, are derived from toxicological studies in animals in which coolants were administered orally ^20^. Calculating MOE for inhalation of a compound based on past oral toxicity studies requires some caution. For human risk assessment and to calculate risk assessment parameters from exposure to these chemicals through various routes, it is routine practice for US and International regulatory agencies (US Environmental Protection Agency, EPA; European Chemical Agency, ECHA; UK’s Interdepartmental Group on Health Risks from Chemicals, IGHRC) to employ route-to-route (R2R) extrapolation ^53,54^. Regulatory agencies agree that the absorption efficiency for chemicals by the respiratory system is higher than the efficiency of the digestive tract and usually, an oral-to-inhalation ratio of 2 is applied for route-to-route extrapolation, reflecting the higher absorption of toxicants following inhalation ^53–55^. In this study, R2R extrapolation was not used, however, if applied, the MOE values would be even lower, reflecting an even higher risk associated with inhalation of these two synthetic cooling agents at various daily consumption volumes. This would suggest even greater urgency for regulatory measures mitigating the risk from WS-3 and WS-23 exposures in E-cigarette users.

Recently, an e-vapor product company (RLX technology, China) has conducted a nose-only acute and sub-acute inhalation study for WS-23 (28-day exposure at doses of 19 and 29 mg/kg-bw/day) ^24^. Whole-body plethysmography was conducted after 28 days of repeated exposure to measure various parameters of pulmonary function. Compared to control and solvent groups, mice exposed to WS-23 demonstrated significant changes in various parameters, such as time of expiratory (Te), peak expiratory flow (PEF), relaxation time (RT), minute volume (MV), respiratory rate (F), end-inspiration pause (EIP), end-expiratory pause (EEP). These results indicate that WS-23 may alter pulmonary function of users of WS-23 containing E-cigarettes. However, the study reports no remarkable histopathological changes in either the respiratory organs (nose, throat, trachea and lungs) or other visceral organs (liver, kidney, heart etc.). It has to be noted that the histopathological analysis was qualitative, and not based on a quantitative histopathologic scoring. The absence of histopathological evidence for organ toxicity contradicts prior WS-23 oral toxicity studies in rats that reported kidney lesions and hepatic toxicity starting at 10 mg/kg-bW/day (NOAEL of 5 mg/kg-bW/day) ^20,21^. The discrepancy between these observations could be due to the different routes of exposure (oral vs. inhalation), or more importantly, may have resulted from the different exposure durations (14-weeks oral dosing vs. 4-weeks inhalational exposure).

*In vitro* genotoxicity studies of WS-23 in mammalian cells showed evidence for clastogenicity ^20^. Clastogenicity was observed only in the presence of metabolic activation, indicating that the effect is mediated by the formation of a reactive metabolite. Due to the increased exposure to WS-23 from use of E-cigarettes, it is imperative that more studies are conducted to understand the mechanisms of their metabolism and further identify WS-23’s reactive clastogenic metabolites.

Previous studies have examined the toxicological effects of synthetic cooling agents, including WS-3 and WS-23, in isolation, and not in mixtures. As demonstrated in this study, refill E-cigarette liquids and popular disposable E-cigarettes contain mixtures of natural and synthetic cooling agents. Inhalation toxicity studies are required to examine the effects of acute, sub-acute and chronic exposures to mixtures of these chemicals and marketed E-cigarettes that contain such mixtures. Another limitation in assessing the risk associated from use of these E-cigarettes is the rapidly evolving product variety and the dynamic nature of their composition. The products analyzed in this study were purchased in the last 1-2 years. More recent marketed products may contain different compositions. Hence, continuous monitoring of new E-cigarette products by determining their chemical compositions for presence of potentially toxic flavorant levels is critical for both regulatory and risk assessment purposes.

Following the withdrawal of flavored Juul cartridges from the US market, disposable E-cigarette products of the brand “Puffbar” brand have become increasingly popular among US youth and young adults ^6,56^. In Puffbar products, synthetic cooling agents were not only found in mint/menthol-flavored, or “cool”- or “ice”-labelled E-liquids, but in almost all products tested (13/14), suggesting that a cooling effect is preferred by consumers. Cooling agents were added to fruit-flavored E-liquids named “Orange Ice” and “Blueberry Ice”, but also “Strawberry Banana” and “Lychee”, without the “Ice” label, and in “Cucumber” and “OMG” flavors. This illustrates the utility of odorless synthetic cooling agents to add a cooling effect to flavors that likely would not be favored by users when combined with menthol, due to the incompatible minty odor the “natural” coolant menthol would add. Synthetic cooling agents may allow tolerance of more intense flavor combinations that would otherwise be irritating.

The fact that synthetic coolants are odorless also raises another important question: Legislation such as the FSPTCA or the European Tobacco Product Directive (TPD) ^1,57^ banned “characterizing” flavorants from combustible tobacco products (with exemption for menthol in the US), yet is unclear how synthetic coolants lacking a characterizing flavor would be treated. It has to be noted that TPD does not allows additives in tobacco products that facilitates inhalation, but no specific compounds that facilitate inhalation or method to measure facilitation are currently listed in the TPD. So, a replacement of the cooling compound menthol (with a minty characterizing flavor) with a synthetic coolants (lacking characterizing flavor) could be a strategy to circumvent a menthol ban. Policy makers should consider this when implementing menthol bans, such as both Canada and Germany have done in respective regulations that either maintain a positive list of compounds permitted as additives in tobacco products (Canada), or specifically ban a whole class of compounds that facilitates inhalation, including all natural and synthetic derivatives of menthol and other cooling agents (Germany) ^58,59^.

## Conclusions

Synthetic cooling agents are added to US-marketed E-cigarettes in a wide range of amounts that are comparable and relative to menthol amounts in flavored E-cigarettes. Synthetic coolants were found not only in mint- and menthol-flavored products, but also in fruit-, dessert- and sweet-flavored E-cigarettes. These coolants are added to refill liquids, E-cigarette liquid diluents and popular disposable E-cigarettes such as Puffbar, at levels that pose potential health risks to users, thereby requiring for these products to be prioritized for risk mitigation measures.

## Supporting information

Jabba_Synthetic Coolant RiskAssessment_Supp555555Data

## Author contributions were as follows

SVJ, HCE and SEJ conceptualized and designed the study; DGT, LD, and HCE acquired and analyzed the chemical analytical data; SVJ analyzed and interpreted the data for risk analysis calculations; SVJ, SEJ and HCE drafted the manuscript; SVJ, HCE, PTA, JBZ, and SEJ provided supervision; JBZ critically revised the manuscript for important intellectual content.

## Data Availability Statement

The data underlying this article are available in the article and in its online supplementary material.

