## Supplementary material for "Synthetic Cooling Agents in US-marketed E-cigarette Refill Liquids and Popular Disposable E-cigarettes: Chemical Analysis and Risk Assessment": Jabba_Synthetic Coolant RiskAssessment_Supp555555Data

3 Genome Ct.

Durham, NC 27710-3094

**Table S1:** Chemical concentrations for menthol, menthone, carvone, synthetic cooling agents (WS-3 and WS-23) and solvent chemicals in US-marketed E-cigarette refill liquids and carryover of each compound. [a]single experiment.

| | | PG/VG ratio | Flavor Concentration ± s.d. (µg/g) [Carryover % ± s.d.] | | | | | Nicotine concentration ± s.d. (mg/g) [Carryover % ± s.d.] |
|---|---|---|---|---|---|---|---|---|
| | **E-cigarette Flavor (Manufacturer)** | | **WS-3** | **WS-23** | **Menthol** | **Menthone** | **Carvone** | |
| 1 | Arctic Rush (Central Vapors) | 50/50 | 14650 ± 230 **[106% ± 4%]** | | | | | 9.2 ± 0.09 **[104% ± 8%]** |
| 2 | Frzn Mango (BLVK) | 30/70 | 2450 ± 00 | | 3640 ± 10 | | | 5.08 ± 0.03 |
| 3 | Nuclear White Menthol (Vapor HQ) | 50/50 | 2440 ± 10 **[104% ± 4%]** | | 23490 ± 130 **[103% ± 2%]** | 310 ± 20 **[106% ± 10%]** | 190 ± 70**[105% ± 7%]** | 9.88 ± 0.02 **[93% ± 4%]** |
| 4 | Rich Girl (Stunna) | 70/30 | 2260 ± 30 **[107% ± 11%]** | | | | | 3.31 ± 0.06 **[94% ± 8%]** |
| 5 | Frozen Berries (Shinjin Vapor) | 30/70 | 2220 ± 20 | | 2320 ± 20 | | | 3.36 ± 0.05 |
| 6 | Frozen Limeade (Shinjin Vapor) | 30/70 | 1540 ± 20 | | 3470 ± 50 | | | 3.38 ± 0.04 |
| 7 | Cool Lime (Chill) | 30/70 | 2110 ± 60 | | | | | 1.63 ± 0.05 |
| 8 | Tropicool (Chill) | 30/70 | 2040 ± 40 **[91%[a]]** | | | | | 3.02 ± 0.07 **[97%[a]]** |
| 9 | Kool Cali (Chill) | 40/60 | 1690 ± 20 | | | | | 4.20 ± 0.06 |
| 10 | Sakura 3 mg Nic (Sugoi) | 30/70 | 1090 ± 20 **[105% ± 2%]** | | 910 ± 10 | | | 2.32 ± 0.03 |
| 11 | Coconut Ice (EC Blend) | 60/40 | 2240 ± 10 | | | | | 1.41 ± 0.02 |
| 12 | Clove + Cool Hit (EC Blend) | 50/50 | 1600 ± 20 | | | | | 17.91 ± 0.23 |
| 13 | Popsicle USA (EC Blend) | 60/40 | 150 ± 10 | | | | 10 ± 1 | |
| 14 | Frozen (FJ's ) | 30/70 | 1550 ± 50 | | | | | |
| 15 | Wintergreen Menthol (VaporHQ) | 50/50 | 870 ± 20 | | 4850 ± 30 | | | 9.72 ± 0.08 |

| | | | | | | | | |
|---|---|---|---|---|---|---|---|---|
| 16 | Icy Menthol (Vista Vapors) | 50/50 | 560 ± 10 | | 9240 ± 90 | 1320± 50 | 2890 ± 10 | 9.80 ± 0.02 |
| 17 | Ice Dragon (ELD Flavors) | 20/80 | 410 ± 30 | | | | | 2.75 ± 0.01 |
| 18 | Wintermint (Gothic Vapor) | 50/50 | 330 ± 50 | | 1110 ± 190 | | 510 ± 10 | 9.03 ± 0.84 |
| 19 | Climate Change (What Juice) | 30/70 | 290 ± 10 | | 330 ± 10 | | | 9.48 ± 0.12 |
| 20 | Arctic (Vista Select) | 30/70 | 280 ± 20 | | 3470 ± 10 | 30 ± 10 | 930 ± 70 | 6.59 ± 0.06 |
| 21 | Dillinger (NastyJuice X Kilo) | 30/70 | | 1860 ± 40 **[99% ± 10%]** | | 280 ± 10 **[85% ± 15%]** | | 4.29 ± 0.07 **[92% ± 11%]** |
| 22 | Kool Peach (Salty Man) | 40/60 | 990 ± 20 **[109% ± 1%]** | | | | | 46.4 ± 1.09 **[109% ± 2%]** |
| 23 | Sakura Nic Salts 35 mg (Sugoi) | 50/50 | 1170 ± 30 | | 1170 ± 20 | | | 28.74 ± 0.43 |
| 24 | Sakura Nic Salts 50 mg (Sugoi) | 50/50 | 1140 ± 60 **[97%[a]]** | | 1140 ± 90 | 14110 ± 670 | | 39.19 ± 2.20 **[94%[a]]** |
| 25 | Unicorn Nicotine Salt Spearmint (BLVK) | 40/60 | 2950 ± 20 **[107% ± 8%]** | | 10920 ± 160 **[107% ± 11%]** | 25450 ± 600 **[88% ± 9%]** | 6830 ± 40 **[97% ± 10%]** | 41.36 ± 0.33 **[106% ± 8%]** |
| 26 | Agent Cool Super Concentrate (TBD Liquids) | | 8840 ± 130 | 124060 ± 720 | | | | |

| | |
|---|---|
| WS-3 | |
| WS-23 | Concentrate |

**Table S2:** Predicted MOE for Synthetic Cooling Agents Containing US-marketed E-Cigarette Liquids Used in Vape Tanks and Vape Mod Devices.

| | E-cigarette Flavor (Manufacturer) | Flavor Concentration (µg/g) | | MOE for Daily E-Liquid (mL/day) Amount-Consumed | | | | |
|---|---|---|---|---|---|---|---|---|
| | | WS3 | WS23 | 1 mL | 3 mL | 5 mL | 10 mL | 15 mL |
| 1 | Arctic Rush (Central Vapors) | 14650 | | 28 | 9 | 6 | 3 | 2 |
| 2 | Frzn Mango (BLVK) | 2450 | | 164 | 55 | 33 | 16 | 11 |
| 3 | Nuclear White Menthol (Vapor HQ) | 2440 | | 171 | 57 | 34 | 17 | 11 |
| 4 | Rich Girl (Stunna) | 2260 | | 192 | 64 | 38 | 19 | 13 |
| 5 | Frozen Berries (Shinjin Vapor) | 2220 | | 181 | 60 | 36 | 18 | 12 |
| 6 | Frozen Limeade (Shinjin Vapor) | 1540 | | 261 | 87 | 52 | 26 | 17 |
| 7 | Cool Lime (Chill) | 2110 | | 190 | 63 | 38 | 19 | 13 |
| 8 | Tropicool (Chill) | 2040 | | 197 | 66 | 39 | 20 | 13 |
| 9 | Kool Cali (Chill) | 1690 | | 242 | 81 | 48 | 24 | 16 |
| 10 | Sakura 3 mg Nic (Sugoi) | 1090 | | 368 | 123 | 74 | 37 | 25 |
| 11 | Coconut Ice (EC Blend) | 2240 | | 190 | 63 | 38 | 19 | 13 |
| 12 | Clove + Cool Hit (EC Blend) | 1600 | | 261 | 87 | 52 | 26 | 17 |
| 13 | Popsicle USA (EC Blend) | 150 | | 2834 | 945 | 567 | 283 | 189 |
| 14 | Frozen (FJ's ) | 1550 | | 259 | 86 | 52 | 26 | 17 |
| 15 | Wintergreen Menthol (VaporHQ) | 870 | | 479 | 160 | 96 | 48 | 32 |
| 16 | Icy Menthol (Vista Vapors) | 690 | | 604 | 201 | 121 | 60 | 40 |
| 17 | Ice Dragon (ELD Flavors) | 410 | | 961 | 320 | 192 | 96 | 64 |
| 18 | Wintermint (Gothic Vapor) | 330 | | 1263 | 421 | 253 | 126 | 84 |
| 19 | Climate Change (What Juice) | 290 | | 1384 | 461 | 277 | 138 | 92 |
| 20 | Arctic (Vista Select) | 280 | | 1433 | 478 | 287 | 143 | 96 |
| 21 | Dillinger (NastyJuice X Kilo) | | 1860 | 135 | 45 | 27 | 13 | 9 |
| 22 | Kool Peach (Salty Man) | 990 | | 413 | 138 | 83 | | |
| 23 | Sakura Nic Salts 35 mg (Sugoi) | 1170 | | 356 | 119 | 71 | | |
| 24 | Sakura Nic Salts 50 mg (Sugoi) | 1140 | | 366 | 122 | 73 | | |
| 25 | Unicorn Nicotine Salt Spearmint (BLVK) | 2950 | | 139 | 46 | 28 | | |
| 26 | Agent Cool Super Concentrate (TBD Liquids) (recommended use at 1-3%) | 8840 (88.4-265.2) | 124060 (1241-3722) | | | | | |
| | 1% Dilution | 88.4 | | 5221 | 1740 | 1044 | 522 | 348 |
| | 3% Dilution | 265.2 | | 1740 | 580 | 348 | 174 | 116 |
| | 1% Dilution | | 1241 | 232 | 77 | 46 | 23 | 15 |
| | 3% Dilution | | 3722 | 78 | 26 | 16 | 8 | 5 |

| | |
|---|---|
| MOE > 100 (Risk Mitigation need not be prioritized) | WS-3 |
| MOE < 100 (Risk Mitigation to be Prioritized) | WS-23 |
| | Concentrate |

**Table S3:** Predicted MOE for Synthetic Cooling Agents Containing Popular Disposable E-Cigarettes, Puffbar.

| | Puffbars Flavors | Flavor Concentration (µg/g) | | WS-3 MOE for Daily E-Liquid Amount Consumed (Puffbars/day) | | | | WS-23 MOE for Daily E-Liquid Amount Consumed (Puffbars/day) | | |
|---|---|---|---|---|---|---|---|---|---|---|
| | | WS-3 | WS-23 | 1/2 Puffbar | 1 Puffbar | 2 Puffbars | | 1/2 Puffbar | 1 Puffbar | 2 Puffbars |
| 1 | Cool Mint | 10770 | 36660 | 60 | 30 | 15 | | 11 | 6 | 3 |
| 2 | Café Latte | | 2300 | | | | | 176 | 88 | 44 |
| 3 | Orange Ice | 1750 | 22950 | 353 | 177 | 88 | | 17 | 8 | 4 |
| 4 | Pomegranate | | | | | | | | | |
| 5 | Lush Ice | | 25830 | | | | | 16 | 8 | 4 |
| 6 | Melon Ice | | 8810 | | | | | 47 | 23 | 12 |
| 7 | Cucumber | 3770 | 3940 | 173 | 87 | 43 | | 104 | 52 | 26 |
| 8 | Lychee | | 26240 | | | | | 15 | 8 | 4 |
| 9 | Menthol | 3540 | 1610 | 179 | 90 | 45 | | 246 | 123 | 62 |
| 10 | Blueberry Ice | 1880 | 14120 | 341 | 170 | 85 | | 28 | 14 | 7 |
| 11 | Strawberry Banana | | 5420 | | | | | 74 | 37 | 18 |
| 12 | OMG | | 7390 | | | | | 54 | 27 | 14 |
| 13 | Peach Ice | | 11540 | | | | | 35 | 18 | 9 |
| 14 | Banana Ice | | 13110 | | | | | 31 | 15 | 8 |

| | |
|---|---|
| MOE > 100 (Risk Mitigation need not be prioritized) | WS-3 |
| MOE < 100 (Risk Mitigation to be Prioritized) | WS-23 |
